# Robertsonian translocations in Danish sika deer (*Cervus nippon*). Markers for absent F1-hybridization with red deer (*C. elaphus*) and implications for selection, speciation and infertility

**DOI:** 10.64898/2026.08.10.743091

**Authors:** Niels Tommerup, Kasper Kristoffer Alsing, Esben Budtz-Jørgensen, Flemming Thune-Stephensen, Astrid Jorun Ingstrup

## Abstract

EU has reclassified the sika deer (*Cervus nippon*) as an undesirable invasive species based on reports that hybridization with the indigenous red deer (*C. elaphus*) may produce fertile offspring. Since sika-derived DNA previosuly introduced into the red deer population (introgression) cannot be removed, the crucial question is whether new (F1) hybridisation occur. To address this, we analysed the chromosomes in 56 sika and 22 red deer. All red deer had a chromosome number 2n=68. In contrast, the chromosome number in sika ranged from 64 to 67, due to the variable presence of two sika-specific Robertsonian translocations (ROB1,ROB2). In the free-ranging sika population in Jutland, >90% of the sika deer were homozygote for at least one of these ROBs, excluding that they could be F1-hybrids. Moreover, ROB2 was in Hardy-Weinberg equilibrium, further supporting the absence of gene flow between the two species. In contrast, ROB1 was in Hardy-Weinberg disequilibrium, suggesting negative fitness of heterozygotes, including potential F1-hybrids. In Jægersborg Deer Park, the eight examined sika deer had the same genotype (absence of ROB1, homozygosity of ROB2), supporting that it is a founder population which may have been isolated for ∼100 years. Again, none of these can be F1-hybrids due to the homozygosity of ROB2. We conclude that F1-hybridisation between sika and red deer either does not occur or occur very rarely in Denmark. The study establish the Danish sika-populations as unique models for adressing important biological questions: What underlies the absence of hybridisation? Why are ROBs frequent in sika deer but not in the closely related red deer? How fast do new species/subspecies develop in isolated founder populations? Which factors determine, that some ROBs have little heterozygous effects, whereas others are selected against, with implications for the role of ROBs as genetic barriers promoting speciation, and for fertility problems in some human ROB carriers.

## Introduction

The present study was initiated following the EU reclassification of the sika deer (*Cervus nippon*) as an undesirable invasive species, primarily based on reports from the UK and Ireland but also continental Europe showing hybridisation with native red deer (Goodman et al., 1999; Herzog & Harrington, 1991; Senn & Pemberton, 2009; Biedrzycka et al., 2012; Putnová et al., 2021). In areas where hybridization has been documented, it occurs in both directions, and in both cases may result in fertile offspring. During the approximately 125 years that sika deer have been present in Denmark (Bennetsen, 1976), no Danish reports of hybrids have occurred.

Any sika-derived DNA introduced into the red deer population through historical hybridization and subsequent backcrossing (introgression) is effectively irreversible and cannot be removed from the population (e.g. Goodman et al, 1999), even if all sika deer would be eradicated. Hence, the crucial question is whether new hybridization (F1-hybridization) regularly occur in Denmark? To assess this, we collected blood samples for chromosome analysis, testes for analysis of chromosome behavior during meiosis, and tissue samples for future DNA analyses. As a first-tier, we choose chromosome analysis since it would allow the detection of Robertsonian translocations (ROB), and secondly, it is a relatively fast and cost-efficient approach.

Genetic studies based on microsatellite and mitochondrial geotyping as well as whole genome sequencing have shown, that sika deer can be divided into two major lineages: the continental Asian and Japanese lineages. The latter can further be divided into Northern and Southern Japanese groups. Both continental and Japanes groups are further divided into several subspecies (Ba et al., 2015; Dong et al., 2022; Liu et al., 2025; Wang et al., 2025).

The chromosome number of Japanese sika deer varies from 64 to 67 (Gustavsson & Sundt, 1968; Omura et al., 1983), whereas the chromosome number in continental Asian sika deer range from 62 to 68 chromosomes (Gustavsson & Sundt, 1969; Mayr & Kalat, 1989). These variations in chromosome number are due to the presence of one or more Robertsonian translocations (ROBs), which are the most common chromosomal rearrangements found in nature. In contrast, red deer have a stable chromosome number of 68 without ROBs, apart from an ancient metacentric chromosome wich is shared with both sika and fallow (*Cervus dama*) deer (Goldoni et al., 1984; Gustavsson & Sundt, 1968; Herzog, 1987b; Proskuryakova et al., 2022).

A ROB is formed when two acrocentric chromosomes fuse into a single meta- or submetacentric chromosome. Variation in chromosome number within a population arises because some individuals carry two copies of a given ROB (= two chromosomes), others carry one ROB together with the two unfused chromosomes involved in the fusion (= three chromosomes), while still others lack the ROB entirely but possess two copies of each unfused acrocentric chromosome (= four chromosomes). Although the total chromosome number differs, all of these combinations result in the same number of chromosome arms (NF = 70) with the same content of unique genetic materiel.

When individuals with and without a ROB interbreed, some offspring will carry a single ROB copy (i.e., be heterozygous). This may reduce fertility or even cause infertility, thereby creating a genetic barrier which have been proposed to contribute to the formation of new species or subspecies (Faria & Navarro, 2010; Galindo et al., 2021; Searle & Hughes, 2025). However, although a ROB is theoretically expected to have a negative fitness effect (i.e., to be underdominant), studies of other wild populations, e.g. mice, have shown that some ROBs are not strongly underdominant but can be in Hardy-Weinberg (HW) equilibrium (Nachman & Myers, 1989). Hardy-Weinberg equilibrium for a sika-specific ROB would indicate that hybridization between sika and red deer cannot be widespread, since one assumption of HW equilibrium is that alleles are neither introduced into nor removed from the population, in this case ROBs from sika entering the red deer population and the corresponding free acrocentric chromosomes from red deer entering the sika population. Moreover, approximately one in ∼1,000 humans carries a ROB (Nielsen & Wohlert, 1991; Therman et al., 1989), most often without any clinical problems. However, for unknown reasons a subset of ROB carriers have fertility issues and/or increased risk of spontaneous miscarriage and liveborn offspring with unbalanced chromosome disorders, e.g. Down and Patau syndrome (trisomy 21 and 13)(for a review, see Jia et al., 2023).

Since the Danish sika population is estimated to be ∼2,100 individuals, approximately 20-25 times smaller than the red deer population (∼50,000 individuals), and because of the reported chromosomal variation in sika (Miyake et al, 1982; Omura et al, 1983; Herzog, 1987a; Mayr & Kalat, 1989), we focused primarily on sika deer. Nevertheless, the study also includes the first cytogenetic study of Danish red deer.

## Results

### Samples

Blood samples were obtained from 80 sika deer and 29 red deer originating from 21 localities (Figure S1; see Materials and Methods). Successful chromosome analyses were obtained from 56 sika deer (70%) and 22 red deer (75.8%). Unsuccessful analyses were primarily due to contamination with bacteria, yeast, fungi, and protists, either individually or as mixed infections. In addition, several blood samples were completely or partially coagulated, most likely because of the interval between culling in the field and blood collection. Overall, more metaphase chromosome spreads were obtained from four-day cultures, and fresh samples (0-1 days old) consistently yielded better results than older samples (up to six days old).

### Chromosome analyses of red deer

All red deer examined, irrespective of locality, exhibited a karyotype consisting of 68 chromosomes, of which chromosome 1 (M1) and the Y chromosome are metacentric, while all remaining chromosomes are acrocentric (Figure 1).

**Figure 1.**
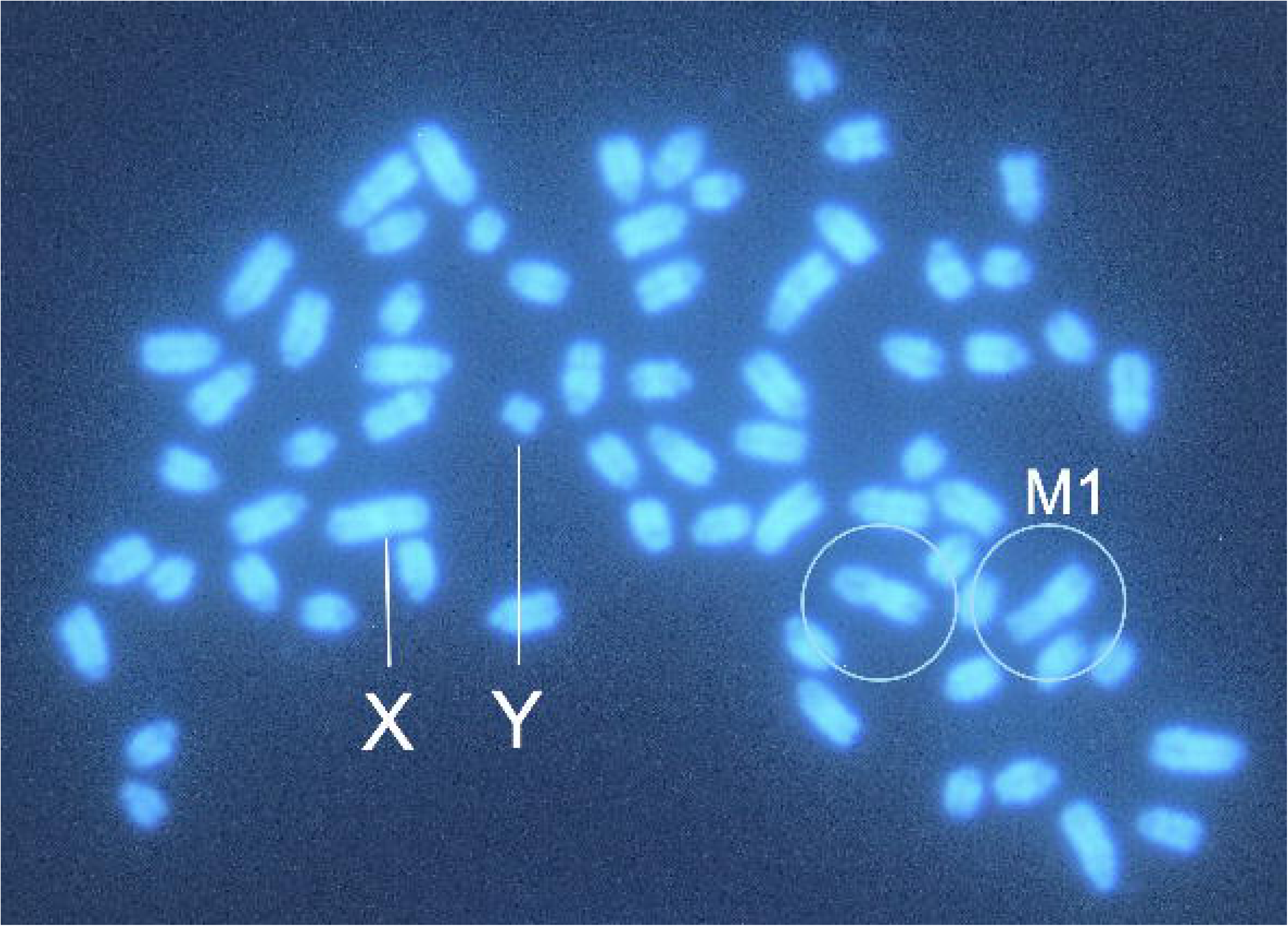
Red deer metaphase spread with 68 chromosomes. Yellow circles on the metacentric chromosome 1 (M1) shared with sika deer.

### Chromosome analyses of sika deer

Three different metacentric autosomes (ROB1, ROB2, and M1) were identified in the sika populations (Figure 2). M1 which was present in two copies in all sika and red deer examined, corresponds to the medium-sized metacentric chromosome previously reported in all investigated sika and red deer, as well as in fallow deer (Mayr et al., 1987). M1 therefore originated before the divergence of *C. nippon* and *C. elaphus* ∼2-3.6 million years ago (Gilbert et al., 2006; Hu et al., 2019). The roe deer (*Capreolus capreolus*) has 70 chromosomes (Rubini & Fontana, 1988) corresponding to the 70 chromosome arms found in red deer, fallow deer, and sika deer. The close evolutionary relationship among these deer species is further reflected by the identical banding patterns observed for the individual chromosomes (Mayr et al., 1987). In red deer, M1 is the largest chromosome and is therefore designated chromosome 1. Since M1 is not informative with respect to the objectives of the present study, it was not included in the subsequent analyses but served as an internal control.

**Figure 2.**
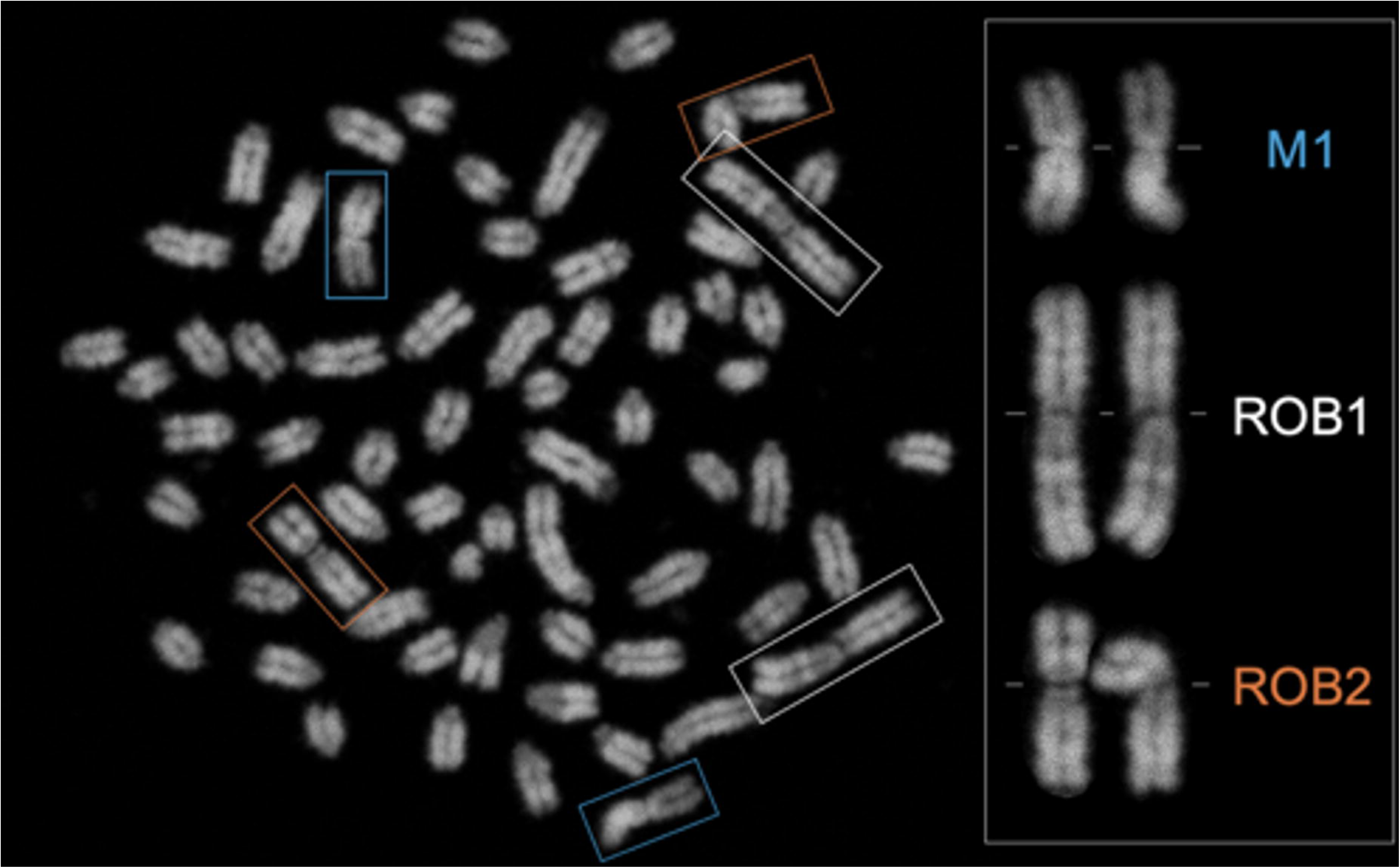
Metaphase spread from a Danish sika stag from Jutland with 64 chromosomes. Two copies of each of the three metacentric chromosomes (M1, ROB1, and ROB2) are present. These chromosomes have been enlarged and arranged on the right. The centromeres, corresponding to the primary constrictions, are indicated by white lines behind the enlarged chromosomes.

Each copy of a Robertsonian translocation present in an individual reduces the diploid chromosome number (2n) by one. Consequently, sika deer lacking both ROB1 and ROB2 would have 68 chromosomes, the same diploid number as red deer. No such animal was identified in the Danish sika population. Sika deer with 67 chromosomes carry a single copy of either ROB1 or ROB2. Individuals with 66 chromosomes possess two ROB copies, which may consist of two copies of ROB1, two copies of ROB2, or one copy of each translocation (Figure 3, bottom row). Deer with 65 chromosomes carry three ROB copies, again in different combinations of ROB1 and ROB2. All individuals with 64 chromosomes possess two copies of both ROB1 and ROB2 (Figure 3)(Table S1).

**Figure 3.**
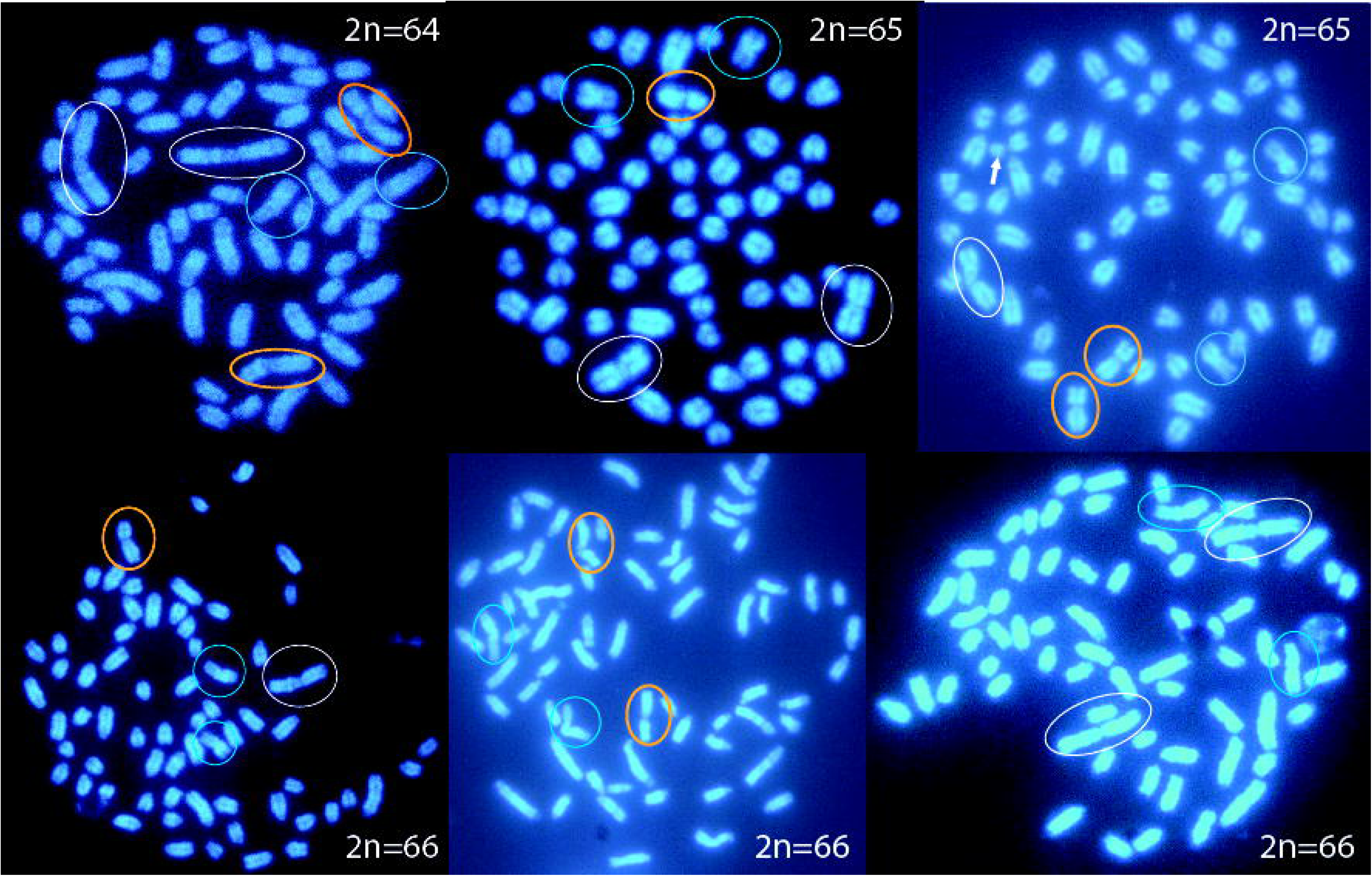
DAPI-stained metaphase spreads from sika deer showing different combinations of the variable Robertsonian translocations ROB1 (white circles) and ROB2 (orange circles). M1 is indicated by blue circles. 2n = diploid chromosome number. The same chromosome number may result from different combinations of ROB1 and ROB2. See also Table 2.

Regardless of the distribution, there are always 70 chromosome arms in both sika deer and red deer (Tables 1 and 2), excluding the Y chromosome in males. The metacentric Y chromosome is the smallest chromosome in male sika deer. It cannot be reliably distinguished from the Y chromosome of red deer and is therefore not informative. Likewise, we disregard the very small short arms of the remaining acrocentric chromosomes, which probably do not carry unique genetic information but consist of C-band positive repetitive DNA (Herzog, 1987b).

**Table 1.**
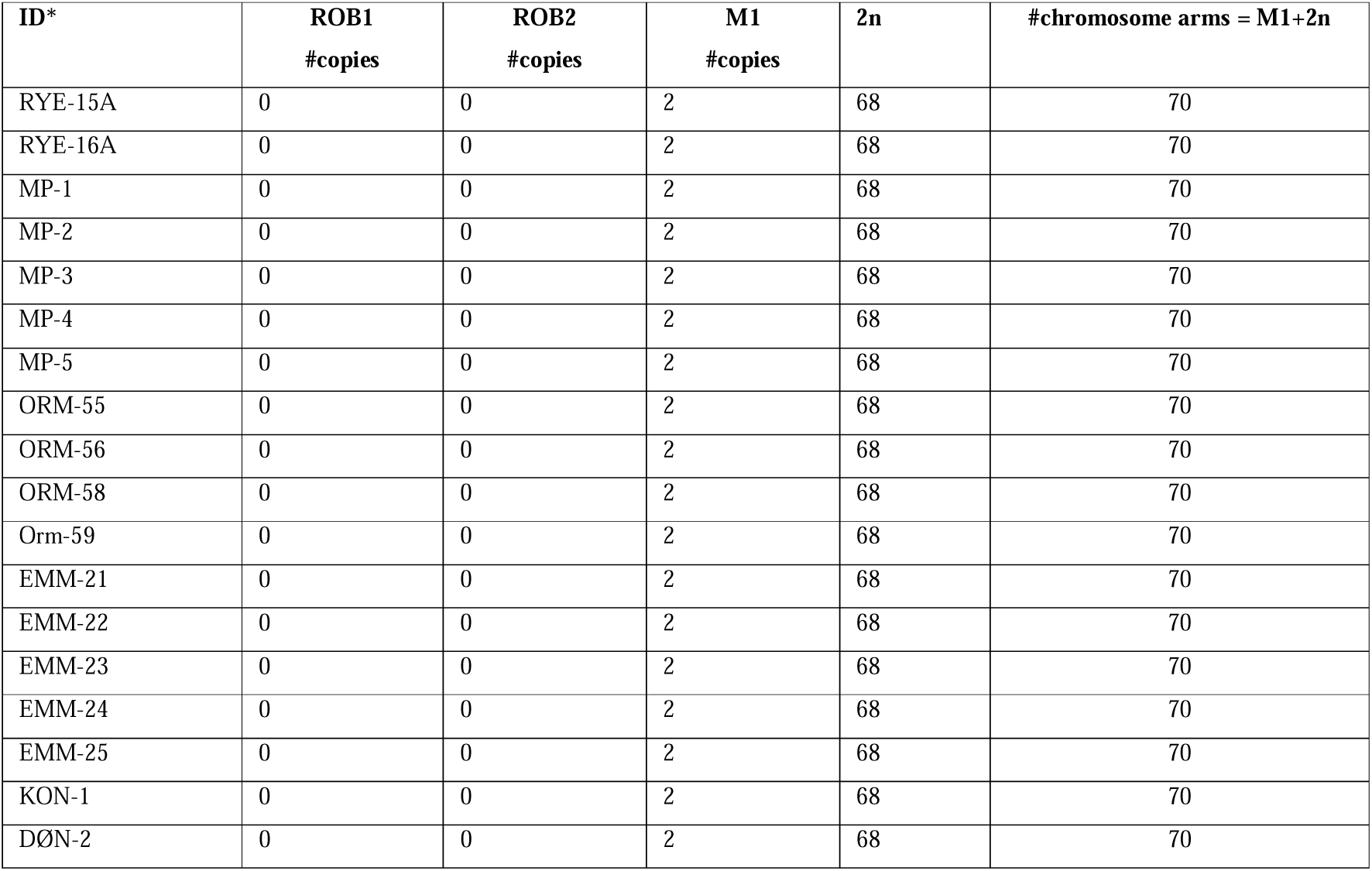
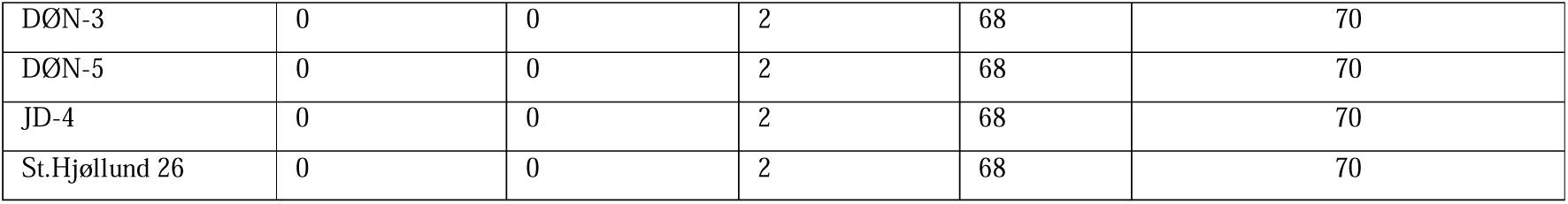
Chromosome analyses of red deer. **ROB1/ROB2** = the sika-specific Robertsonian translocations, which were not detected in red deer (0). **2n** = diploid chromosome number (number of chromosomes in somatic cells).

**Table 2.**
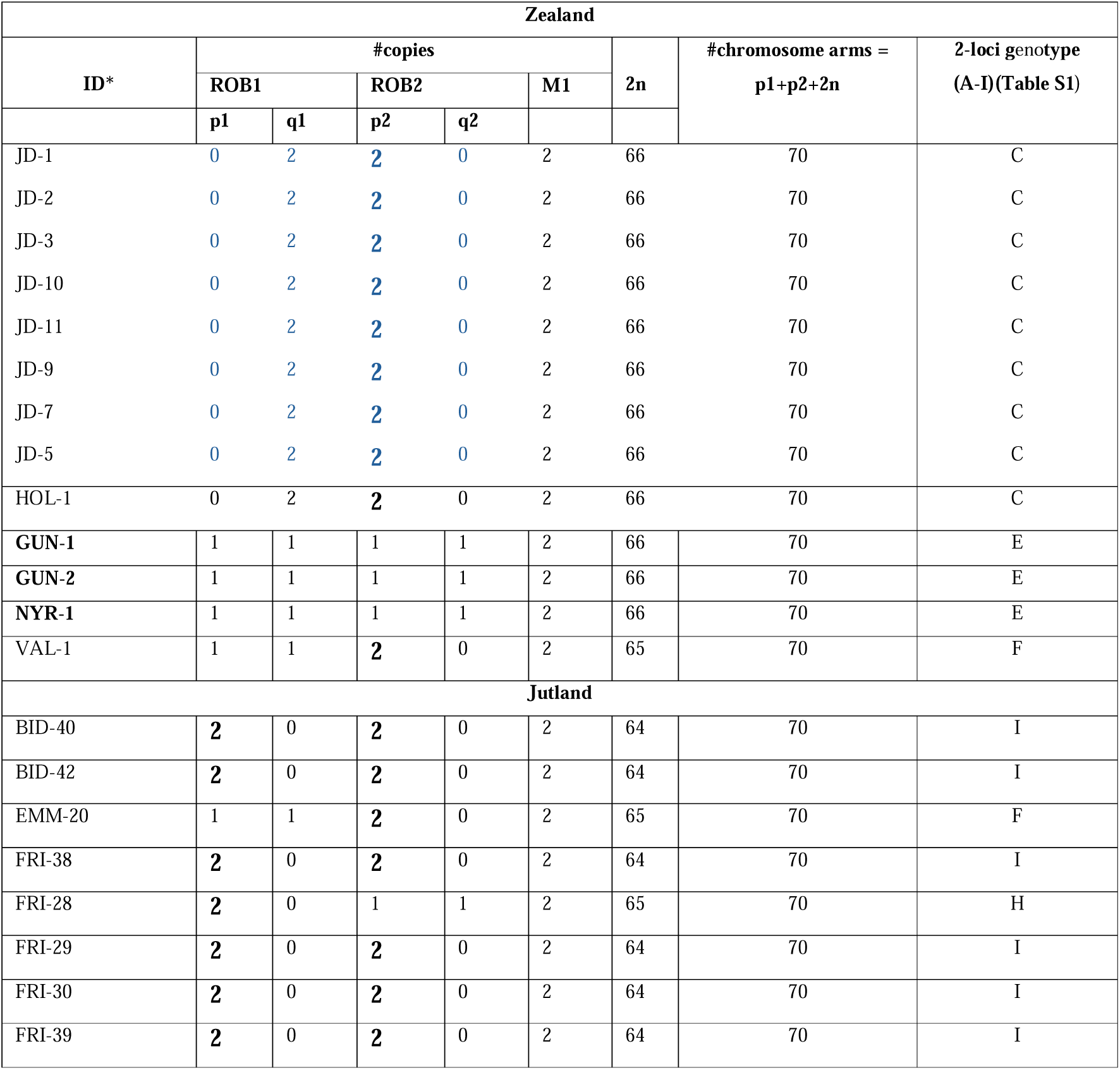
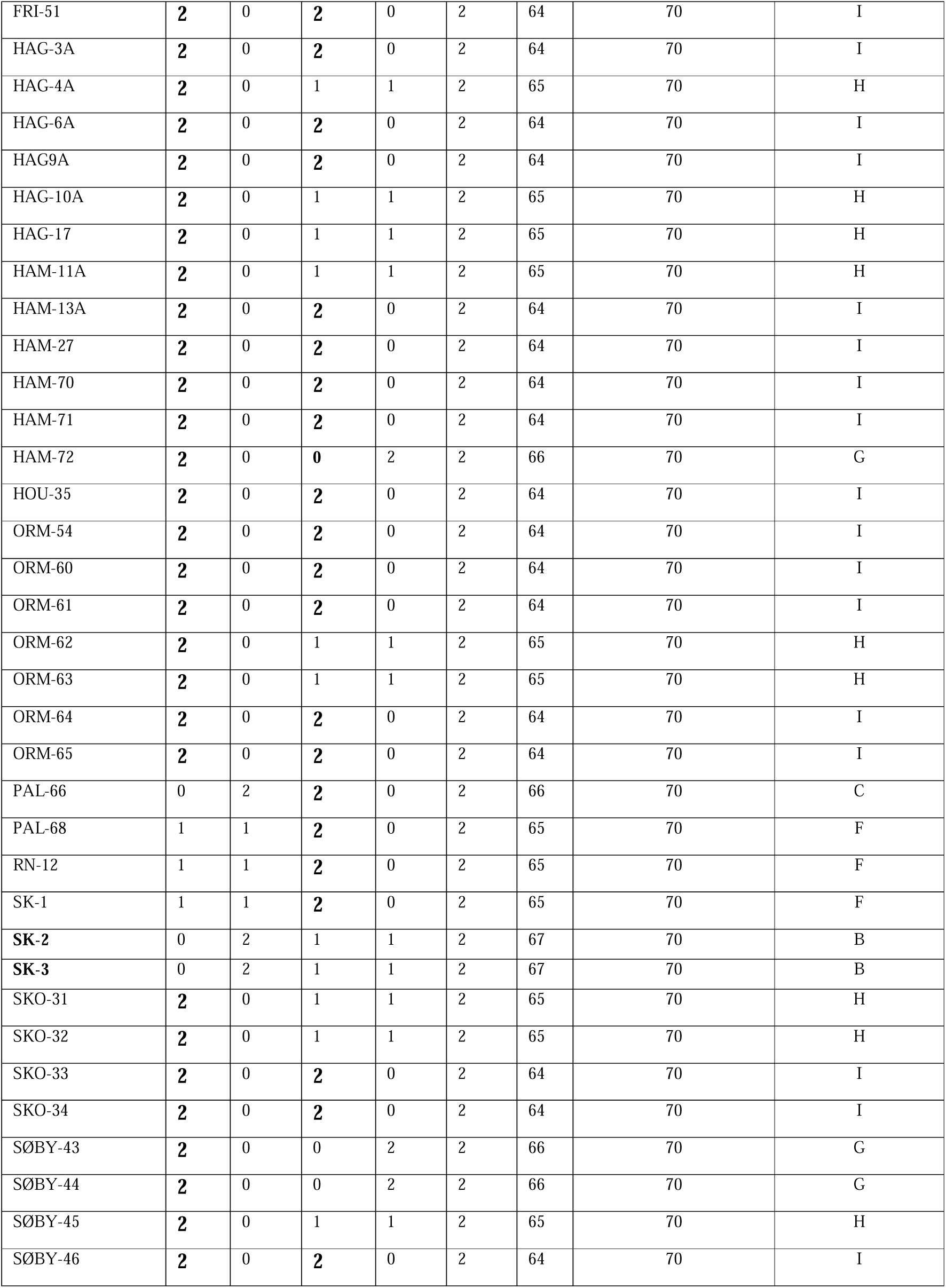
Chromosome analyses of sika deer. ROB1/ROB2 = the sika-specific Robertsonian translocations. p denotes the number of copies of a given ROB. q denotes the number of copies of the two free chromosome arms involved in the specific ROB. **Larger font with bold type** indicate homozygosity for a ROB. ID with **bold type** indicates animals that are not homozygous for at least one ROB.

| Zealand |  |  |  |  |  |  |  |  |
| --- | --- | --- | --- | --- | --- | --- | --- | --- |
| ID* | #copies |  |  |  |  | 2n | #chromosome arms =<br>p1+p2+2n | 2-loci genotype<br>(A-I)(Table S1) |
|  | ROB1 |  | ROB2 |  | M1 |  |  |  |
|  | p1 | q1 | p2 | q2 |  |  |  |  |
| JD-1 | 0 | 2 | 2 | 0 | 2 | 66 | 70 | C |
| JD-2 | 0 | 2 | 2 | 0 | 2 | 66 | 70 | C |
| JD-3 | 0 | 2 | 2 | 0 | 2 | 66 | 70 | C |
| JD-10 | 0 | 2 | 2 | 0 | 2 | 66 | 70 | C |
| JD-11 | 0 | 2 | 2 | 0 | 2 | 66 | 70 | C |
| JD-9 | 0 | 2 | 2 | 0 | 2 | 66 | 70 | C |
| JD-7 | 0 | 2 | 2 | 0 | 2 | 66 | 70 | C |
| JD-5 | 0 | 2 | 2 | 0 | 2 | 66 | 70 | C |
| HOL-1 | 0 | 2 | 2 | 0 | 2 | 66 | 70 | C |
| GUN-1 | 1 | 1 | 1 | 1 | 2 | 66 | 70 | E |
| GUN-2 | 1 | 1 | 1 | 1 | 2 | 66 | 70 | E |
| NYR-1 | 1 | 1 | 1 | 1 | 2 | 66 | 70 | E |
| VAL-1 | 1 | 1 | 2 | 0 | 2 | 65 | 70 | F |
| Jutland |  |  |  |  |  |  |  |  |
| BID-40 | 2 | 0 | 2 | 0 | 2 | 64 | 70 | I |
| BID-42 | 2 | 0 | 2 | 0 | 2 | 64 | 70 | I |
| EMM-20 | 1 | 1 | 2 | 0 | 2 | 65 | 70 | F |
| FRI-38 | 2 | 0 | 2 | 0 | 2 | 64 | 70 | I |
| FRI-28 | 2 | 0 | 1 | 1 | 2 | 65 | 70 | H |
| FRI-29 | 2 | 0 | 2 | 0 | 2 | 64 | 70 | I |
| FRI-30 | 2 | 0 | 2 | 0 | 2 | 64 | 70 | I |
| FRI-39 | 2 | 0 | 2 | 0 | 2 | 64 | 70 | I |
| FRI-51 | 2 | 0 | 2 | 0 | 2 | 64 | 70 | I |
| HAG-3A | 2 | 0 | 2 | 0 | 2 | 64 | 70 | I |
| HAG-4A | 2 | 0 | 1 | 1 | 2 | 65 | 70 | H |
| HAG-6A | 2 | 0 | 2 | 0 | 2 | 64 | 70 | I |
| HAG9A | 2 | 0 | 2 | 0 | 2 | 64 | 70 | I |
| HAG-10A | 2 | 0 | 1 | 1 | 2 | 65 | 70 | H |
| HAG-17 | 2 | 0 | 1 | 1 | 2 | 65 | 70 | H |
| HAM-11A | 2 | 0 | 1 | 1 | 2 | 65 | 70 | H |
| HAM-13A | 2 | 0 | 2 | 0 | 2 | 64 | 70 | I |
| HAM-27 | 2 | 0 | 2 | 0 | 2 | 64 | 70 | I |
| HAM-70 | 2 | 0 | 2 | 0 | 2 | 64 | 70 | I |
| HAM-71 | 2 | 0 | 2 | 0 | 2 | 64 | 70 | I |
| HAM-72 | 2 | 0 | 0 | 2 | 2 | 66 | 70 | G |
| HOU-35 | 2 | 0 | 2 | 0 | 2 | 64 | 70 | I |
| ORM-54 | 2 | 0 | 2 | 0 | 2 | 64 | 70 | I |
| ORM-60 | 2 | 0 | 2 | 0 | 2 | 64 | 70 | I |
| ORM-61 | 2 | 0 | 2 | 0 | 2 | 64 | 70 | I |
| ORM-62 | 2 | 0 | 1 | 1 | 2 | 65 | 70 | H |
| ORM-63 | 2 | 0 | 1 | 1 | 2 | 65 | 70 | H |
| ORM-64 | 2 | 0 | 2 | 0 | 2 | 64 | 70 | I |
| ORM-65 | 2 | 0 | 2 | 0 | 2 | 64 | 70 | I |
| PAL-66 | 0 | 2 | 2 | 0 | 2 | 66 | 70 | C |
| PAL-68 | 1 | 1 | 2 | 0 | 2 | 65 | 70 | F |
| RN-12 | 1 | 1 | 2 | 0 | 2 | 65 | 70 | F |
| SK-1 | 1 | 1 | 2 | 0 | 2 | 65 | 70 | F |
| SK-2 | 0 | 2 | 1 | 1 | 2 | 67 | 70 | B |
| SK-3 | 0 | 2 | 1 | 1 | 2 | 67 | 70 | B |
| SKO-31 | 2 | 0 | 1 | 1 | 2 | 65 | 70 | H |
| SKO-32 | 2 | 0 | 1 | 1 | 2 | 65 | 70 | H |
| SKO-33 | 2 | 0 | 2 | 0 | 2 | 64 | 70 | I |
| SKO-34 | 2 | 0 | 2 | 0 | 2 | 64 | 70 | I |
| SØBY-43 | 2 | 0 | 0 | 2 | 2 | 66 | 70 | G |
| SØBY-44 | 2 | 0 | 0 | 2 | 2 | 66 | 70 | G |
| SØBY-45 | 2 | 0 | 1 | 1 | 2 | 65 | 70 | H |
| SØBY-46 | 2 | 0 | 2 | 0 | 2 | 64 | 70 | I |

### F1-hybridization?

In Jutland, more than 95% (41/43) of the examined sika deer have two copies of either ROB1 and/or ROB2 (i.e., they are homozygous) (highlighted with larger & bold font in Table 2). None of these can therefore be F1-hybrids with red deer, since red deer lacks these ROBs. At the same time, any potential red deer/sika F1-hybrid would have at least one of these two ROBs. The remaining 2 out of 43 (<5%) sika deer are heterozygous for ROB2; hence, half of their potential F1-hybrid offspring would also inherit one copy of ROB2. The absence of sika-specific ROBs among the examined red deer therefore makes it highly unlikely that any of the investigated red deer are F1-hybrids.

On Zealand, two sika genotypes were primarily identified. In Jægersborg Deer Park, all examined sika deer (8/8) share the same genotype for the two ROBs (lacking ROB1 and carrying two copies of ROB2). This genotype was found in only one of the 43 sika deer examined in Jutland. The difference is highly significant (100% vs 2,3%, p < .001, Fisher’s exact test). The marked overrepresentation of this genotype in Jægersborg Deer Park compared with Jutland support that the Jægersborg population originated from a small, isolated founder population. Further support for this is, that the same genotype was found in the one examined sika from Holbæk Deer Park, which received its sika deer from Jægersborg Deer Park 40-50 years ago (Torben Christiansen, personal communication). Apart from excluding F1-hybrids in the examined sika deer in Jægersborg (and Holbæk) Deer Parks, any F1-hybrid among the red deer would carry one copy of ROB2.

The second Zealand genotype - with one copy of ROB1 and one copy of ROB2 (i.e. double heterozygotic) - was found in the three examined sika deer from southern Zealand: two from Gunderslevholm and one from nearby Nyrup Hegn. The latter animal most likely escaped from Gunderslevholm. In theory, the examined sika deer from Gunderslevholm could have been F1-hybrids with a red deer. However, the sampled animals were a calf and a young hind. Since the last red deer stag at Gunderslevholm was shot three years before the samples were collected, this excludes the possibility that they are F1-hybrids. Furthermore, the remaining red deer hinds at Gunderslevholm did not become pregnant by the remaining sika stags, despite the fact that hinds normally conceive every year.

Among the nine possible genotype combinations of ROB1 and ROB2, seven have been identified in Denmark (Table 2; Table S1).

### Hardy-Weinberg analysis of the sika population in Jutland

The largest free-ranging population of sika deer is found in East/Central Jutland, which was also among the first regions where sika deer were introduced into Denmark around 1900. The animals originated from Hagenbeck in Hamburg, possibly via Sweden (Bennetsen, 1976). It is unclear whether they were pure *C. nippon nippon* or belonged to one or more other subspecies of *C. nippon*.

Being free-ranging, the Jutland sika population has had the opportunity to disperse among nearby forests during the more than 100 years since its introduction. Consequently, the samples from Jutland can be treated as originating from a single population inhabiting one large, connected forest area with unrestricted movement and random mating - one of the prerequisites for Hardy-Weinberg equilibrium.

For ROB1, 36 individuals carried two copies, 4 individuals carried one copy, and 3 individuals lacked ROB1 entirely. This distribution shows a significant difference between the observed and expected genotype frequencies (*p*-value: 0.0003), indicating that **ROB1 is not** in Hardy-Weinberg equilibrium within the population (Figure 4A,C). For ROB2, 28 individuals carried two copies, 12 individuals carried one copy, and 3 individuals lacked ROB2. This distribution yielded a *p*-value of 0.30, indicating no significant difference between the observed and expected genotype frequencies. ROB2 is therefore in Hardy-Weinberg equilibrium within the population (Figure 4B,C).

**Figure 4.**
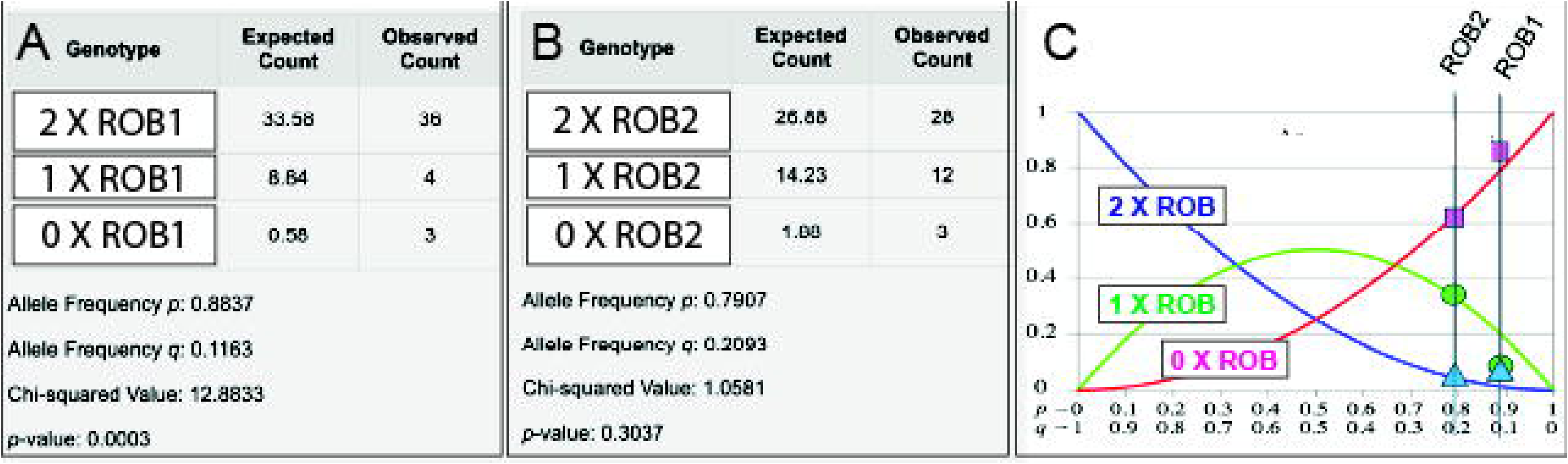
Hardy-Weinberg calculations and visualization of Hardy-Weinberg equilibrium for ROB2 and Hardy-Weinberg disequilibrium for ROB1, the latter showing an excess of homozygotes and a deficit of heterozygotes. Images are modified from https://ug.sebc.me/labs/hwe-calculator.

If we exclude the most geographically separated sample in Jutland (EMM-20), originating from the Djursland peninsula (Figure S1), having one copy of ROB1 and two copies of ROB2)(Table 2), this does not change the HW-conclusions for ROB1 (*p*-value: < 0.0001) and ROB2 (*p*-value: 0.3261).

## Discussion

This first Danish chromosome analysis of sika deer and red deer is also one of the largest cytogenetic studies of its kind in the world. We show, that F1-hybridization between sika and red deer either does not occur in Denmark, or if it does occur, it must be extremely rare. We confirm that Danish red deer possess 68 chromosomes, as reported in all previous studies. We identify two sika-specific Robertsonian translocations, ROB1 and ROB2, where more than 91% of Danish sika deer are homozygous for at least one of these translocations, excluding that they are F1-hybrids. Chromosome analysis is therefore a reasonably reliable and inexpensive method for screening for F1-hybridization in Denmark. It may also be a necessary method, since Robertsonian translocations cannot be detected by DNA sequencing nor by any other methods previously used for assessing hybridization (e.g. microsatellite analysis, mitochondrial genotyping). We show that ROB2 is in Hardy-Weinberg equilibrium in the Jutland population, providing further evidence that there is no significant gene flow between sika deer and red deer. We also show that ROB1 is not in Hardy-Weinberg equilibrium, suggesting possible selection against heterozygous carriers, including potential F1-hybrids.

It is unknown which subspecies of sika deer have been introduced into Denmark, but they are reportedly descended from temple deer originating from the southern Japanese islands where they are considered sacred animals. Whether ROB1 and ROB2 are identical to previously reported Robertsonian translocations in sika deer is uncertain, due to the different staining/banding techniques employed, from conventionally stained, R-banded, C-banded, and G-banded chromosomes (Fontana & Rubini, 1990; Gustavsson & Sundt, 1969; Herzog, 1987a, 1987c; Miyake et al., 1982; Proskuryakova et al., 2022; Herzog & Harrington, 1991a) to our use of DAPI-banding. However, ROB1 and ROB2 resembles those reported in sika deer morphologically classified as *C. nippon nippon* (Herzog, 1987a), which would be in line with a southern Japanese origin.

So far, there does not appear to be a straightforward relationship between karyotype and individual subspecies. However, only a few of the many recognized sika subspecies (Ba et al., 2015; Møller-Nielsen, 2026) have been cytogenetically investigated. Based on size and centromere index other Robertsonian translocations occur in sika deer in addition to those identified in the present study. This includes an early study of Japanese sika deer, where the chromosome numbers ranged from 64 to 67 chromosomes (Omura et al., 1983). Robertsonian translocations are the most common chromosomal rearrangement in nature, and most newly arising translocations are likely eliminated rapidly through selection. A general assumption is that once homozygosity (fixation) has been established, a Robertsonian translocation may function as a species-specific reproductive barrier because backcrossing results in reduced fertility, e.g. due to frequent unbalanced chromosome segregation during gametogenesis (Galindo et al., 2021; Gerton, 2024; Schubert, 2024). In this context, it is noteworthy that homozygosity for ROB1 predominates in the Jutland sika population, while homozygosity for ROB2 is likely present in the entire sika population in Jægersborg Deer Park. In theory, this high degree of homozygocity could represent relative barriers against hybridization with red deer and thus provide a possible genetic explanation for the apparent absence of hybridization in Denmark. Another fascinating possibility is that the apparent fixation of ROB2 observed in Jægersborg Deer Park may facilitate the emergence of a new sika subspecies. We do not know how quickly species or subspecies arise, but the approximately 100-year history of a likely founder lineage in Jægersborg Deer Park offers a unique opportunity to investigate this.

Although Robertsonian translocations are proposed to be primary drivers of chromosomal evolution (and thus speciation), and although they may be associated with infertility, subfertility, chromosomal imbalance, and disease (for example, in humans and cattle), there are many examples in nature where Robertsonian translocations are in Hardy-Weinberg equilibrium (e.g., in mice), while others are not (Searle & Hughes, 2025). The observed Hardy-Weinberg disequilibrium of ROB1 in the Jutland sika population may represent a Robertsonian translocation in the process of becoming fixed, whereas ROB2 appears to have a minor negative effect enabling it to persist as a polymorphism in Hardy-Weinberg equilibrium.

Two hypotheses have been proposed to explain the origin of Robertsonian translocations in deer. The first suggests that hybridization itself initiates Robertsonian translocations (Herzog & Harrington, 1991a). One of their arguments was, that all continental sika deer were hybrids between red deer and sika deer. This hypothesis is not supported by recent DNA data, which clearly distinguish Japanese sika from continental forms. Also, the finding of chromosome number variation (2n = 64-68) due to Robertsonian translocations in wild sika populations from different regions of Japan (Omura et al., 1983) contradicts this conclusion. An alternative explanation is that different sika subspecies in geographically isolated regions of Japan possessed distinct Robertsonian translocations, which subsequently became mixed through inter-subspecific hybridization within hybrid zones. The second hypothesis proposes that Robertsonian translocations arise spontaneously and may then spread rapidly - or even become fixed - in small populations with high levels of inbreeding, depending on selection and random genetic drift (Neitzel, 1982). Our study support the this hypothesis.

One intriguing question is why sika deer exhibit a diversity of Robertsonian translocations, whereas red deer apparently never do. The first detailed characterization of human Robertsonian translocations using long-read sequencing technologies and telomere-to-telomere annotated assemblies may help answer this question in the future (de Lima et al., 2025; Mostovoy et al., 2024). Perhaps, like humans, sika deer possess inverted repetitive sequences around the centromeres that facilitate recombination and the formation of Robertsonian translocations (de Lima et al., 2025).

The absence of F1-hybridization in Denmark is consistent with other evidence. During the approximately 125 years that sika deer have been present in Denmark, hunters and wildlife managers have not reported hybrid animals, despite the fact that hybrids elsewhere frequently exhibit altered body size, skull morphology, and antler characteristics (Groves, 2006; Senn et al., 2010). Moreover, DNA analyses of two sika deer and one red deer from Jægersborg Deer Park, examined in Poland as part of a European hybridization study (Biedrzycka et al., 2012), were ultimately excluded from publication because they proved to be pure representatives of their respective species (Aleksandra Biedrzycka, personal communication). Moreover, even in regions where hybridization has been most extensively documented (i.e. Scotland), it is a rare event and extreme hybridization is confined to small geographical areas (Senn & Pemberton, 2009).

One of the important questions in evolutionary biology is, to what extent and how chromosomal rearrangements including ROBs may act as reproductive barriers contributing to speciation. One the one hand, ROBs in Hardy-Weinberg equilibrium in wild populations show that not all ROBs are strong reproductive barrier candidates, e.g. Nachman & Myers, 1989; Faria & Navarro, 2010; Galindo et al, 2021; Searle & Hughes, 2025. In this context the lack of ROB1 heterozygotes establish the sika population in Jutland as a genotyped, large mammalian and naturally occuring model, where the sampled testes can be used to study the meiotic disturbancies proposed to underly ROB-associated negative fitnes. These mechanisms include abnormal chromosome pairing (delayed synapsis/asynapsis), abnormal chromosomal segregation leading to increased rates of aneuploidy (Bonnet-Garnier et al., 2006, 2008), abnormal meiotic recombination, and silencing due to association with the XY-body. For reviews of proposed models and mechanism underlying the chromosomal theory of speciation, see Faria & Navarro, 2010 and Garagna et al, 2014.

Furthermore, the karyotyped cohort can be used for future whole genome sequencing to illuminate the unique 125-year history of the Danish sika population and enable a comprehensive genetic analysis of populations and individuals with known Robertsonian translocations. The elegance of this lies in the possibility of monitoring these processes over generations, incorporating state-of-the-art molecular analyses of both individual animals and entire populations, including studies of gametogenesis in carriers with the different combinations of Robertsonian translocations. Whereas hybridization between Scottish red deer and sika deer has become a model system for studying hybridization, Danish sika populations may become models for investigating why hybridization does not occur and why different Robertsonian translocations are or are not selected against. The implications for our understanding of the role of ROBs in speciation, and for infertility, subfertility and chromosomal disorders in humans are obvious.

## Materials and Methods

Before the 2025 winter culling, sample kits containing tubes for blod sampling (4.5 ml Venoject tubes), muscle tissue (10 ml tubes prefillede with 70% ethanol) and testes (50 ml Falcon tubed prefilled with 30 ml methanol:glacial acetic acid (3:1)) and a sterile Petri dish were sent to Flemming Thune-Stephensen and Astrid Jorun Ingstrup, who took most of the samples. The blood samples were collected in the field under as sterile conditions as possible and as quickly as possible after shooting (ranging from minutes to ∼2 hours), by exposing and sampling from the jugular veins. A muscle biopsy from the same region was placed in the tube containing alcohol. Testes were cleaned of hair and membranes and cut into approximately 0.5-1 cm transverse slices in a Petri dish. The slices were transferred to the 50 ml Falcon centrifuge tube containing 3:1-fixative. Samples were kept refrigerated and transferred to the laboratory within a few days. Muscle samples were drained of alcohol, divided into two portions, and stored at −20°C for future DNA analyses. Liekwise, testes preserved in 3:1-fixative are stored in a refrigerator for future analyses.

All blood samples (n = 109) were cultured under sterile conditions using short-term culture methods. Approximately 0.5-0.6 ml whole blood was added to 7.5 ml PB-MAX Karyotype Medium (Gibco) containing antibiotics in upright T25 tissue culture flasks with filter caps (Nunc EasYFlask 25 cm², Nunclon Delta Surface, vented cap, Thermo Scientific). After 3-4 days of incubation at 37°C in a 5% CO atmosphere, KaryoMAX Colcemid (Gibco) (0.5 μg/ml) was added for 1 hour. Following centrifugation, the cell pellet was treated with 8 ml of 0.075 M hypotonic KCl solution for 30 minutes, centrifuged again, and fixed under agitation with freshly prepared 3:1 methanol:glacial acetic acid. The fixation procedure was repeated three times.

Microscope slides were cleaned overnight in 99% ethanol, rinsed seven times in Milli-Q water, and stored refrigerated in Milli-Q water. Ten microliters of fixed cell suspension were dropped onto a cold, wet microscope slide and air dried. Three drops of VECTASHIELD Antifade Mounting Medium with DAPI (Vector Laboratories) were applied and covered with a 24 × 60 mm coverslip. After sealing with nail polish and drying, preparations were scanned with a fluorescence microscope (Leica DMR). Suitable metaphases were photographed with a 63× oil-immersion objective and a DeltaPix camera. Between three and ten informative metaphase spreads from each animal were analyzed for chromosome number and for the presence or absence of metacentric chromosomes/Robertsonian translocations.

## Ethics and Legislation

All samples were taken on dead animals during the regular winter culling, in compliance with applicable Danish hunting legislation, specifically the *Danish Hunting and Wildlife Management Act* (https://www.retsinformation.dk/eli/lta/2023/639), the *Statutory Order on Open Seasons for Certain Mammals and Birds* (https://www.retsinformation.dk/eli/lta/2024/470), and the *Statutory Order on the Release of Game, Hunting Methods, and Hunting Equipment* (https://www.retsinformation.dk/eli/lta/2017/1652).

## Statistics

For Hardy-Weiberg calculations and graphs we used the Seb Carvallo Hardy-Weinberg Equilibrium Calculator (https://ug.sebc.me/labs/hwe-calculator). Fisher Exact test was conducted in R 4.4.3 to compare genotype frequencies in Jægersborg Deer Park vs. Jutland.

## Contributions

AJS, NT and FT-S designed the study. AJS and FT-S collected and shipped the samples to NT. NT, and KKA cultured, harvested and fixed the blood samples, prepared the slides and analysed the chromosomes. EBJ did statistical analyses. NT interpreted the results and wrote the first draft manuscript. Everybody read and commented on the first draft and on the final manuscript.

## Aknowledgements

This study was supported by the Danish Forest Association (DFA) who had no influence on the results nor the interpretation of the results. Annemette Friis Mikkelsen and Rabab Rima are thanked for their expert technical help. Merete Fredholm is thanked for reading and commenting on the final manuscript.

