## Supplementary material for "Robertsonian translocations in Danish sika deer (*Cervus nippon*). Markers for absent F1-hybridization with red deer (*C. elaphus*) and implications for selection, speciation and infertility": Robertsonian translocations in Danish sika deer_Supplementary.docx

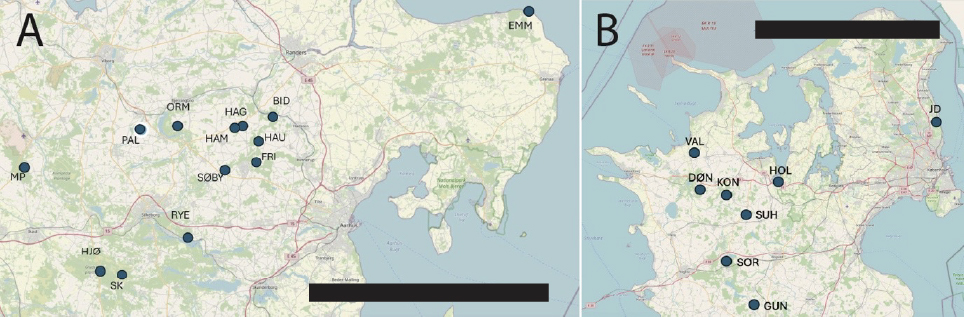

***Figure S1.*** *Geographical origin of the samples with associated forest areas:* ***A) Jutland:*** *SK:Skærbæk Plantation. FRI:Frijsenborg Estate. ORM:Ormstrup Estate. SKO:Skovfryd. Søby:Søbygård. RYE:Rye Nørskov Estate; BID:Bidstrup Estate. HAM:Hammel Forest. HAG:Hagsholm. EMM:Emmedsbo Plantation. HAU:Haurum Forest. PAL:Palstrup Estate. MP:Myremalm plantation.* ***B) Zealand.*** *DØN:Dønnerup Estate. HOL:Holbæk Deer Park. VAL:Valdemarskilde Estate. KON:Kongsdal Estate. GUN:Gunderslevholm Estate. SUH:Den Suhrske stiftelse. SOR:Sorø Forests. JD:Jægersborg Deer Park. Bars = 50 km.*

***Table S1****. The nine possible genotypes (A-I) of ROB1 and ROB2. The number of copies of the specific ROB og the corresponding free acrocentric chromosomes are indicated with 0 (no ROB and two copies of the free chromosomes; 1 ROB and therefore also 1 copy of the free chromosomes (heterozygotic); or 2 ROB copies (homozygotic) and thus no free chromosomes. Note that the same number of chromosomes (2n) can be due to different combinations of ROB1 and ROB2. p1 and p2 = ROB1 and ROB2, respectively. q1 and q2: The free chromosomes involved in ROB1 and ROB2, respectively. See also Figure 3.*

|  | **ROB1** | | **ROB2** | |  |  |
| --- | --- | --- | --- | --- | --- | --- |
| **Type** | **p1** | **q1** | **p2** | **q2** | **Number of observed animals** | **Brief explanation** |
| **A** | 0 | 2 | 0 | 2 | 0 | No ROB1, no ROB2 (2n=68) |
| **B** | 0 | 2 | 1 | 1 | 2 | No ROB1, one ROB2 (2n=67) |
| **C** | 0 | 2 | 2 | 0 | 10 | No ROB1, two ROB2 (2n=66) |
| **D** | 1 | 1 | 0 | 2 | 0 | One ROB1, no ROB2 (2n=67) |
| **E** | 1 | 1 | 1 | 1 | 3 | One ROB1, one ROB2 (2n=66) |
| **F** | 1 | 1 | 2 | 0 | 5 | One ROB1, two ROB2 (2n=65) |
| **G** | 2 | 0 | 0 | 2 | 3 | Two ROB1, no ROB2 (2n=66) |
| **H** | 2 | 0 | 1 | 1 | 10 | Two ROB1, one ROB2 (2n=65) |
| **I** | 2 | 0 | 2 | 0 | 23 | Two ROB1, two ROB2 (2n=64) |
|  |  |  |  |  | **56** |  |
